# Rock Buntings in Central Europe: Phylogeographic connectivity and a current inventory in the Upper Middle Rhine valley

**DOI:** 10.1101/2025.03.20.644321

**Authors:** Michael Wink, Hedwig Sauer-Gürth, Hermann Willems, Ingolf Schuphan

**Affiliations:** Universität Heidelberg; Institute of Pharmacy and Molecular Biotechnology (IPMB), Im Neuenheimer Feld 364, D-69120 Heidelberg; Hermann Willems, Department of Veterinary Clinical Sciences, Clinic for Swine, Justus Liebig University Giessen, Frankfurter Strasse 114, D-35392 Giessen; Institute for Plant Physiology (Bio III), Aachen University (RWTH), Worringerweg 1, D-52074 Aachen

**Keywords:** Phylogeography, microsatellites, cytochrome b, connectivity, gene flow, population decline

## Abstract

The Rock Bunting (*Emberiza cia*) occurs from the Iberian Peninsula to mountainous regions in Central Asia. In Europe, its centre of distribution lays in the Mediterranean region. A small subpopulation is found in Germany in rocky habitats (often with vineyards) along the rivers Middle Rhine, Nahe, Main, Moselle and Ahr. In this study, the German population of the Rock Bunting was monitored for more than 15 years and a continuous decline was observed. Buntings were mist netted and colour ringed; furthermore, blood samples were collected for a DNA analysis. For 122 samples the variability of nucleotide sequences of the mitochondrial cytochrome b gene and the allele distribution of 12 polymorphic microsatellite markers were investigated. Both mtDNA and nuclear DNA show genetical variability, but no geographic clustering, indicating a connectivity and gene flow between the German subpopulations.

## 1. Introduction

The Rock Bunting (*Emberiza cia*) inhabits the temperate and Mediterranean zones, rocky steppe and mountainous regions of the southern Palaearctic and Central Asia (Glutz von Blotzheim and Bauer 1997; Keller et al. 2020) (Fig. 1). In Central Europe, the species is mainly distributed in the montane zone (Schmid et al. 1998; Maumary 2007) up to 2300 m above sea level in open rocky and scrubby habitats (Schuphan and Wink 2016; Schuphan 2011b).

**Figure 1:**
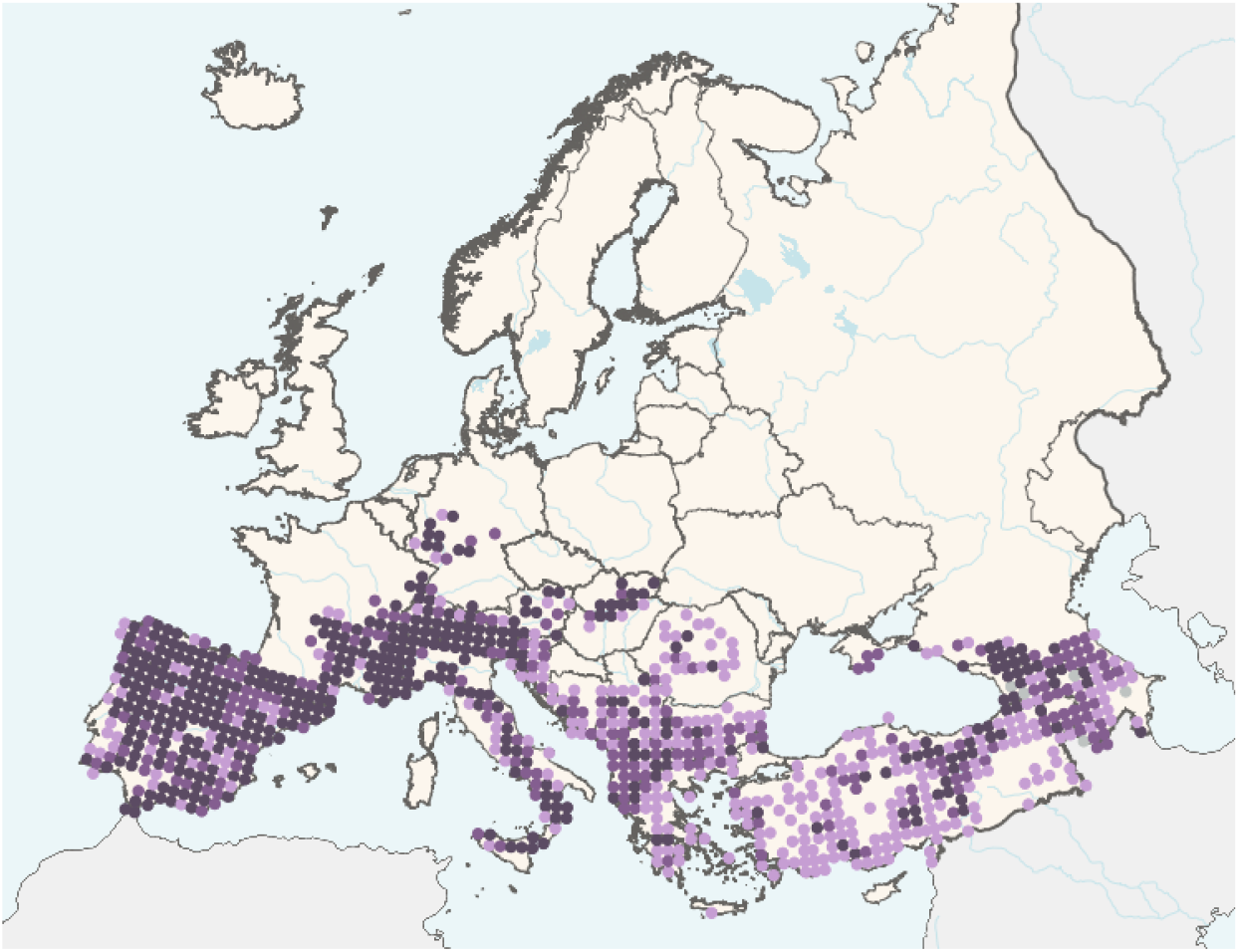
Distribution of the Rock Bunting in Europe (from EBBA 2; https://ebba2.info)

In its northernmost distribution areas in Central Europe, the Rock Bunting is a rare breeding bird. It is particularly widespread in climatically favoured areas along the river slopes of the Rhine, Main, Nahe, Moselle and Ahr in xerothermic rocky habitats including vineyard terraces (Schuphan 2011a, 2017). A recent decline of the Rock Buntings was recorded in some mountains in Germany (Black Forest, Palatinate) which was probably due to the loss of typical habitats (steep clearings, former rocky mountain pastures). These Rock Buntings did not move to the vineyards adjacent to the hillsides, as evident in the Palatinate (Schuphan 2011b).

A population of Rock Buntings in the Upper Middle Rhine Valley has been monitored for more than 50 years (Schuphan 1972, 2011a; Schuphan and Flehmig 2022). A decline in Rock Buntings in the years 1960-80, which can be attributed to reorganization processes in the vineyards (Schuphan 2007; Schuphan and Flehmig 2022), drew our interest to the possible exchanges between the buntings of the Rhine valley with those of its tributaries Ahr, Moselle, Nahe, and Main (Fuchs and Macke 2002; Bosselmann 2008; Schlotmann and Dietrich 2012; Schuphan, 2011c; Schuphan 2017) and the isolated mountainous Rock Bunting occurrences of the Northern and Southern Black Forest, Palatinate, Vosges and Canton of Valais (CH) (Dorka 2010; Deuschle 2010; Groh 1988; Pfeffer and Gilot 2002; Keusch 1991; Schmid et al. 1998; Maumary, 2007; Schuphan and Wink 2016) (see Figure 2).

**Figure 2:**
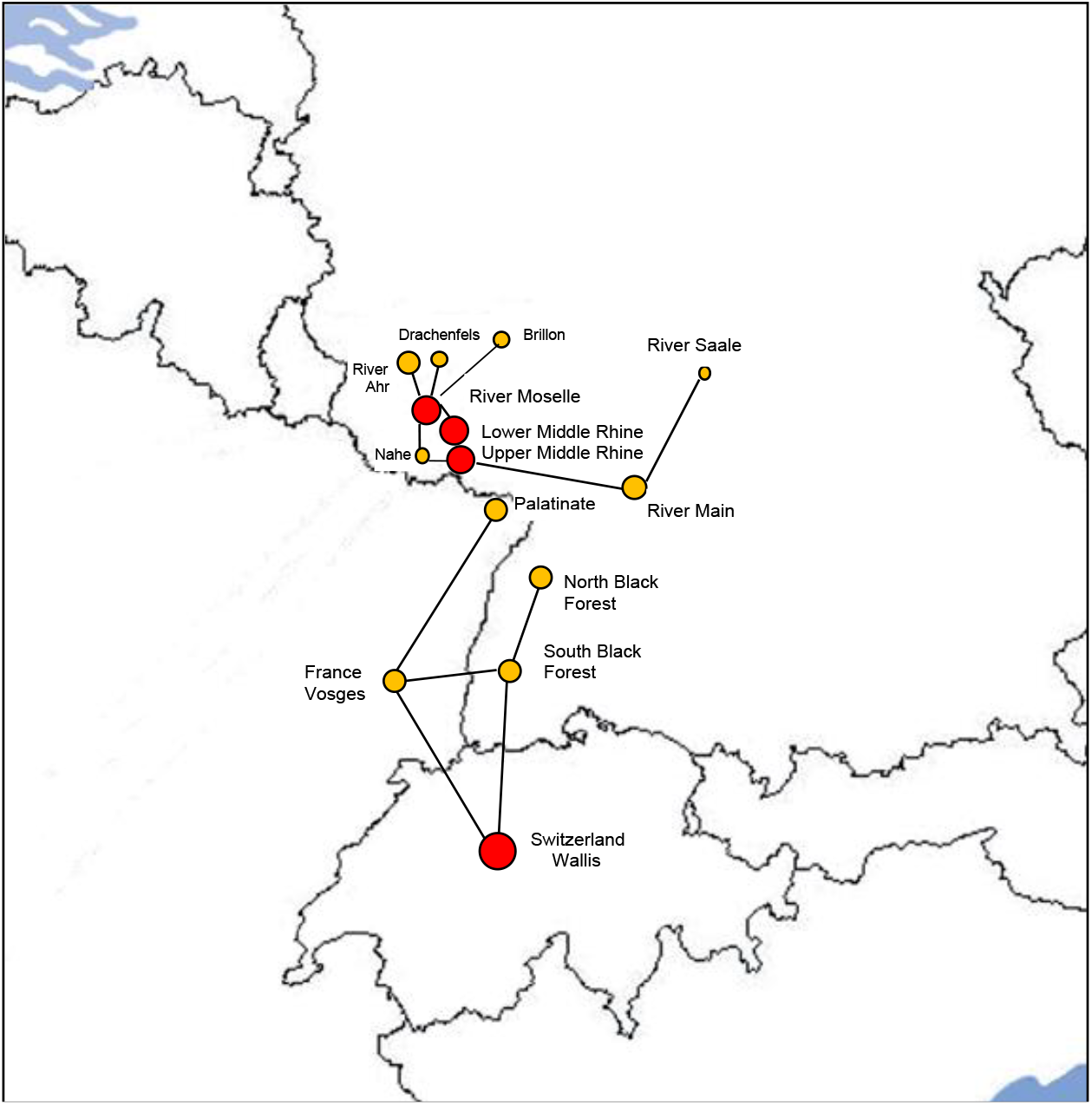
Possible network of Rock Bunting subpopulations in Germany, eastern France and Swiss. red=Rhine- and Canton of Valais-subpopulations = refugium-, colonization centre; ochre = Ahr-; Moselle-; Nahe-; Drachenfels-; Brillon-; Main-; Palatine-, Saale-; North- and South-Black Forest-; Vosges-Subpopulation

A monitoring of all these fragmented habitat patches, combined with partly systematic recordings and colour ringing, led to the impression that these occupied patchily distributed habitats form a network in which the local populations belong to one or two metapopulations as described by the stochastic area occupancy models by Hanski and Ovaskainen (2003). The obvious loss of the Rock Bunting in the Black Forest (Dorka 2009 in Deuschle et al. 2010), in Palatinate (Groh 1988; Schuphan and Grimm 2012) and the general decline in the remaining areas in the last years (Schuphan 2017; Schuphan and Flehmig 2022) supported the assumption of a network between these Rock Bunting habitat patches. It seemed likely that a subpopulation-metapopulation network exists between these habitat patches.

The metapopulation concept describes a population, which includes many spatially discrete subpopulations (Levins 1970). These are linked by dispersal events. The occupied habitat patches depend on the ratio of colonization and extinction rates. For stability of the metapopulation colonization must exceed extinction. In the case of a North- and South-Black Forest and Palatinate subpopulation, extinction could have exceeded, while colonization holds off. The other - vineyard confined Rock Buntings - are probably connected in a network between the habitat patches of the Middle Rhine and its tributary streams Ahr, Moselle, Nahe, Main and form a second metapopulation (Figure 2).

In this communication, we document the results of a monitoring program and have analysed the phylogeographic structure of Rock Buntings from Germany, and a few samples from France and the Iberian Peninsula. We have employed nucleotide sequences of the mitochondrial cytochrome b gene and microsatellite analyses to infer the phylogeography of the Rock Bunting and of its metapopulation in Central Europe.

## 2. Material und Methods

### 2.1 Monitoring and sampling of Rock Buntings

In the Upper Middle Rhine valley (Hesse), the Rock Buntings were systematically monitored and many of them colour-ringed for decades by one of the authors (IS). To get more information of all other subpopulations the relevant regions were visited several times each year from 2007 to 2024, combined with partly systematic recordings, mist-netting and colour ringing (Table S1).

The Rock Buntings of the tributary streams of the Upper Middle Rhine, the Ahr, Moselle, Nahe, and Main inhabit similar steep habitats like at the Rhine near Rüdesheim. Breeding territories are distributed exclusively in southwards directed steep river valley slopes, often planted with vineyards. Usual these habitats form narrow sloop stripes of 100-200 m width between the river and the upper coppice. These slopes are accessible to farmers by restricted roads and these can be used to record the Rock Buntings.

The habitats of the Rock Buntings in the southern mountain areas - the Southern Black Forest, Vosges and Canton of Valais - are completely different from those of the northern vineyard areas in Germany. The breeding sites are characterized as steep rocky, windy, partly cold southerly exposed slopes up to 1300 m above sea level. In the Canton of Valais, the breeding area reaches even 2300 m. In former days, the Palatinate breeding area was characterized as southerly exposure, often even steep forest clearings or rolled lumber areas up till 600 m.

For both types of habitats, play back of the song, the tape lure (TL) were necessary to detect Rock Buntings. It was used repeatedly all 200-300 m in the typical habitats. These are sunny places on rocky slopes in hills and high mountains, ravines, and in clearings of conifer forests. The TL was presented - if possible - mainly using an external car-linked speaker, because the power of the hand-linked TL was often not sufficient. The Rock Bunting males showed up mainly immediately and came close, a few of them watched from far <u>without</u> attracting <u>attention</u> and a few of them sneak up secretly first. After a second back play – at least after 5 min - they all showed up nearby. The documentation of the route and the monitored buntings was performed with a Garmin GPSmap76C receiver and documented using the software MapSource. The records were later processed via Google Earth (details in Schuphan 2011a,b,c). Usually, the territorially reacting males were captured and colour-ringed. All records in this paper refer on territorially reacting Rock Buntings.

### 2.2 DNA analysis

In the context of this project, blood samples (50 – 100 µl) were collected from birds captured for ringing and to examine the phylogeographic network between the Rock Bunting subpopulations.

The blood samples were kept in EDTA buffer (Arctander 1988, Storch et al. 2013) and kept below 20°C in the field, and finally were stored below -18°C. DNA-isolation was performed by the standard phenol-chloroform protocol (Sambrook et al. 1989).

The mitochondrial cytochrome b (Cyt b) gene was amplified by PCR using the following primers: Phoe-L14915: ATGGCCCTCAAYCTHCGTAAAAACC and MT-Fr-s: CAGTTTTTGGTTTACAAGAC. PCR conditions: 94 °C denaturation for 5 min, followed by 39 PCR cycles: 94 °C denaturation 45 sec, 50 °C annealing for 1 min, and 72 °C for extension (2 min). The reaction was executed in a final volume of 50 µl, each reaction containing 100 ng DNA, 2 µL dNTP (each 2.5 mM), PCR Buffer (10x) with 1.5 mM MgCl_2,_ 5 pmol of forward and reverse primers and 1 unit Taq Polymerase.

The PCR products were purified by precipitation in 4 M NH_4_Ac and absolute ethanol (1:1:10), centrifuged for 13,000 rpm for 30 min, followed by centrifugation in 70% ethanol using the same settings and later dissolved in 25 µl sterile H_2_O. For sequencing 1.0 µl sequencing primer (10 pmol/µl) was combined with 7.0 µl PCR products and water. Sequencing primers: Phoe-L14915: ATGGCCCTCAAYCTHCGTAAAAACC and MT-C2: TGAGGAC AAATATCATTCTGAGG. The Sanger sequencing was executed on an ABI 3730 automated capillary sequencer (Applied Biosystems, Carlsbad, CA, USA) with the ABI Prism Big Dye Terminator Cycle Sequencing Ready Reaction Kit 3.1 (carried out by STARSEQ GmbH, Mainz, Sequencing Germany).

The nucleotide sequences were aligned using the program Bio Edit 7.2.5 and WinEdt 5.3. Additionally, we refined the alignment by eye and checked for stop codons in the sequences to exclude the possibility that nuclear copies were amplified. Sequences were analysed by MEGA 11 (Tamura et al., 2021). Sequence polymorphisms and haplotype abundance were calculated with DnaSPv6 (Rozas et al., 2017). A haplotype Network was established using Network 5.0.1.1 from Fluxus-Engeneering.com

For the analysis of nuclear markers, we employed a total of 12 polymorphic microsatellites loci in three PCR panels (Segelbacher and Schuphan 2014) (Table 1). Twelve polymorphic microsatellite loci were genotyped in 122 specimens of the Rock Bunting using the primers shown in Table 1. The Type-it PCR Kit for Microsatellites (Qiagen, Cat. No. 206243, Hilden, Germany) was employed. PCR was carried out in 15 µl reaction volume consisting of 30 ng of total DNA, 0.2-0.3 pmol/µl of each of the forward and reverse primer, 7.5 µl of Master Mix (included in the kit) and a variable amount of RNase-free water to reach the final volume. Conditions for PCR: 95 °C for 5 min, 29 PCR cycles of 95 °C denaturation for 30 sec, 58 °C annealing temperature for 1 min, 72 °C for 30 sec of extension, 60°C for 30 min and 4 °C for 10 min.

**Table 1:**
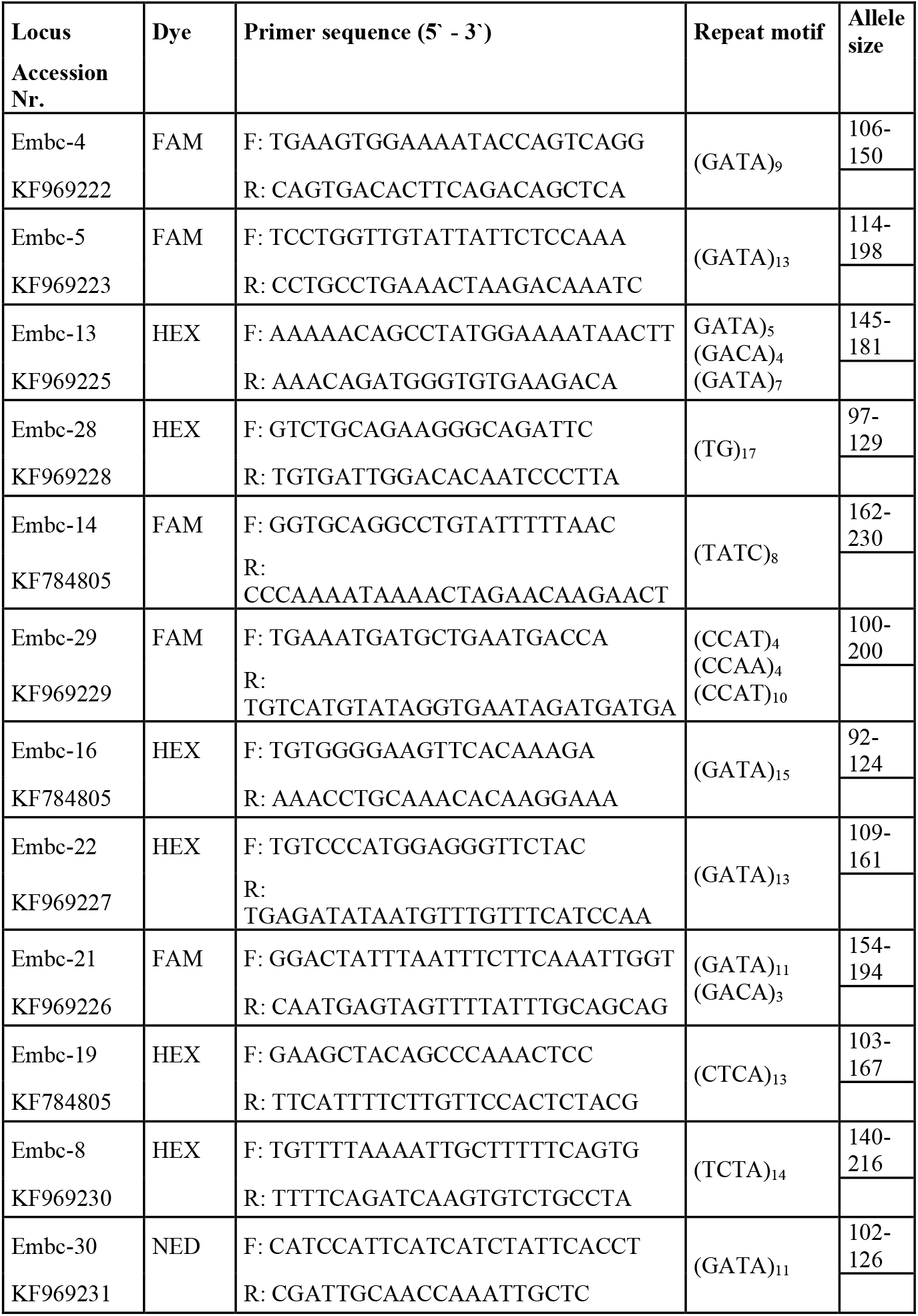
Characterisation of the microsatellite loci analysed in this study.

PCR products were separated by capillary electrophoresis conducted by GATC Biotech AG. The software PEAK Scanner^TM^ version 2.0 was used to determine the allele sizes, in comparison with in-lane size standards (GeneScan^TM^ 500 ROX^TM^, ABI, Cat. No. 401734). We checked for the presence of null alleles and genotyping mistakes (amplification mistakes, recording of stutter peaks and short allele dominance) using the software MICROCHECKER version 2.2.3.

Population genetic analyses were performed mostly within the statistical software R (R Core team, 2017). The function *null*.*all* implemented in the R package PopGenReport v3.0.4 was used to calculate frequencies of null alleles. Deviations from the Hardy-Weinberg equilibrium (HWE) were tested with the function *hw*.*test* implemented in the R package pegas v0.12. The test was performed as an exact test based on Monte Carlo permutations (n=1000) of alleles (Guo and Thompson, 1992).

We employed the function *test_LD* of the R package genepop v1.1.7 for an exact test for genotypic linkage disequilibrium. The length of the dememorization step of the Markov chain was set to 10,000, the number of batches to 1,000, and the iterations per batch to 10,000. Private alleles were determined with functions implemented in the R package poppr v2.8.3. The function *divBasic* implemented in the R package diveRsity v.1.9.90 was used to calculate population genetic parameters (mean number of alleles, rarefied allelic richness, observed heterozygosity, expected heterozygosity, inbreeding coefficient Fis).

The genetic structure was investigated with a Bayesian clustering method and implemented in STRUCTURE version 2.3.4 using the admixture model without prior information about the populations (Pritchard et al. 2000). MCMC parameters were fixed at 50000 and 20000, with run length of 10^6^ iterations and K varying from 1 to 10 clusters. The calculations were repeated five times to confirm convergence among estimated settings. The most probable number of clusters (K) was based on the harmonic mean estimator proposed by Pritchard et al. (2000) and on the second order rate of variation of the likelihood function referring to *K* (Δ*K*) as proposed by Evanno et al. (2005).

Discriminant Analysis of Principal Components (DAPC), implemented in the R package adegenet v2.0.1, was applied as a second approach to detect population structure (Jombart et al. 2010). The number of retained principal components was validated with the cross validation function *xvalDapc*.

## 3. Results and Discussion

### 3.1 Inventory of the of the Rock Bunting in the Upper Middle Rhine

The inventory of the territorial Rock Buntings in the Upper Middle Rhine valley between Lorchhausen and Rüdesheim with 50 territorial males since the start of the monitoring in 1983 is documented in Table 2 and Figure 3. During the years 1960-70 the vineyards of the Upper Middle Rhine valley were reorganized. Small very steep vineyard terraces with their dry masonry walls were removed and larger vineyards were established to facilitate the work in the vineyards and to improve their productivity. These measures lead to a first decline in the population size of the Rock Bunting. In the following years, many steep and not remediated wine terraces, constituting important habitat patches for the buntings, were given up by the farmers and became overgrown; as a consequence, these sites were lost as habitat for the Rock Bunting (Schuphan 2007). A similar trend (albeit hardly documented) was recorded in all isolated populations of the species in Germany (Schuphan 2017).

**Table 2:**
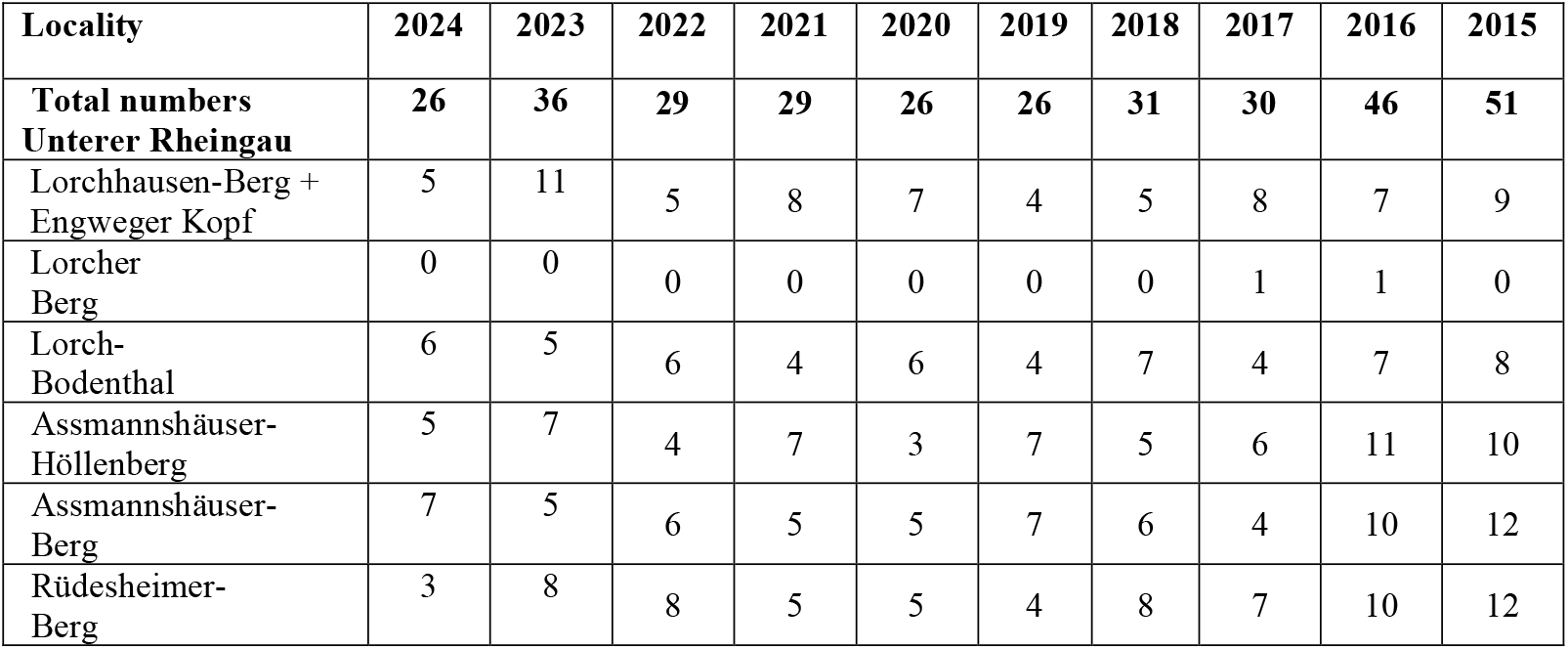
Rock Bunting counts between Lorchhausen and Rüdesheim from 2015 till 2024.

| Locality | 2024 | 2023 | 2022 | 2021 | 2020 | 2019 | 2018 | 2017 | 2016 | 2015 |
| --- | --- | --- | --- | --- | --- | --- | --- | --- | --- | --- |
| **Total numbers**<br>**Unterer Rheingau** | **26** | **36** | **29** | **29** | **26** | **26** | **31** | **30** | **46** | **51** |
| Lorchhausen-Berg +<br>Engweger Kopf | 5 | 11 | 5 | 8 | 7 | 4 | 5 | 8 | 7 | 9 |
| Lorcher<br>Berg | 0 | 0 | 0 | 0 | 0 | 0 | 0 | 1 | 1 | 0 |
| Lorch-Bodenthal | 6 | 5 | 6 | 4 | 6 | 4 | 7 | 4 | 7 | 8 |
| Assmannshäuser-Höllenberg | 5 | 7 | 4 | 7 | 3 | 7 | 5 | 6 | 11 | 10 |
| Assmannshäuser-Berg | 7 | 5 | 6 | 5 | 5 | 7 | 6 | 4 | 10 | 12 |
| Rüdesheimer-Berg | 3 | 8 | 8 | 5 | 5 | 4 | 8 | 7 | 10 | 12 |

**Figure 3:**
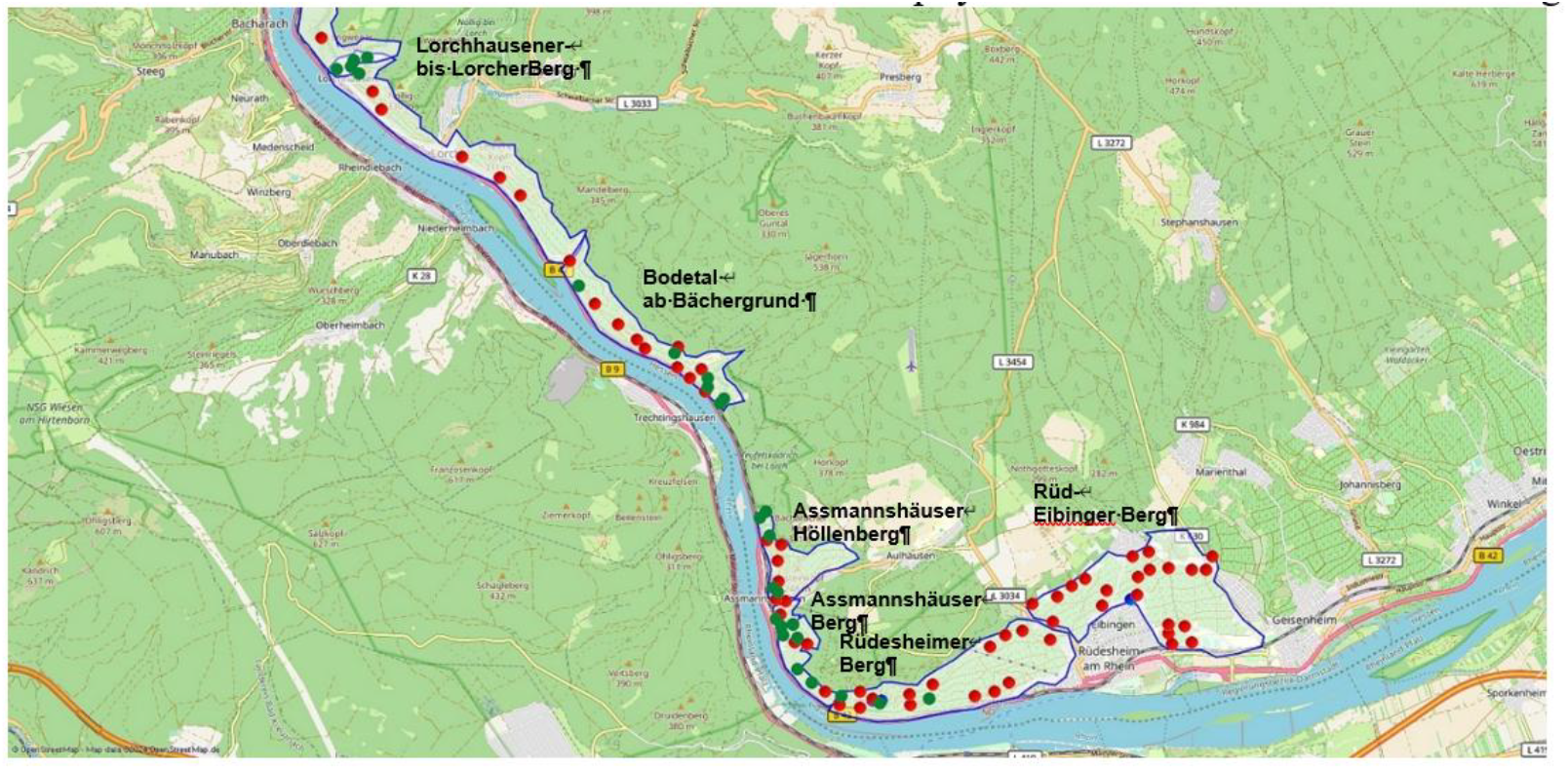
Distribution of the Rock Bunting (green dots) in 2024 Lorchhausen and Rüdesheim (Lower Rheingau) together with Cirl Bunting (red dots) (*E. cirlus*) which has shown a substantial range expansion recently.

Until recently, the total population of Rock Bunting in the Upper Middle Rhine valley between Lorchhausen and Rüdesheim decreased abruptly from about 50 to 26 Rock Buntings in 2024 (Table 2 and Figure 3). Interestingly, another Mediterranean Bunting, the Cirl Bunting (*Emberiza cirlus*) has extended its distribution in a northerly direction in Germany during the last 30 years. It is breeding on slopes of the Rhine rift valley and increasingly in Rock Bunting habitats of the Middle Rhine (Figure 3).

Over 90% of the German population of Rock Buntings live on the slopes of the Middle Rhine and its tributary streams. We distinguish isolated subpopulations at the rivers Ahr, Moselle, Middle-Rhine, Nahe, and the Main (Schuphan 2011c, 2017). In the last ten years, the Ahr subpopulation consisted of about 40 territories, the Moselle subpopulation of about 60 – 80 territories, the Lower Middle Rhine subpopulation of about 50 – 60 territories, the Nahe subpopulation of about 30 territories (Schuphan 2017, Figure 2), the Upper Middle Rhine subpopulation in 2015-2024 of about 50 territories (Table 2), and last, the Main subpopulation, mainly located downstream between Veitshöchheim - Karlstadt (Kalbenstein) and at Homburg (Kallmuth) in 2011 of 32 territories (Schuphan 2011c).

The decline from about 50 to about 30 Rock Buntings in 2017 at the Rhine in its exclusive occurrence in the federal state Hesse, is obviously correlated with wet and cold weather conditions during the first offspring in the preceding year 2016. The February was about + 2.3 °C warmer than average (30-year annual mean). The birds were stimulated too early for breeding. Then a cold-wet March and April period followed. The temperature averaged - 0.7°C for both months below the longterm mean values. April-Mai both reached than 152% of the mean annual rainfall. These weather conditions were negative for the parent birds and their offspring but also for the development of the insect larvae at the border of the coppice, which developed two weeks too late. The loss of Rock Buntings was not compensated in 2018 (Table 2).

### 3.2 Phylogeography of Rock Buntings in Germany

#### 3.2.1 Analysis of cyt b sequences

In a first set of analyses, we sequenced the mitochondrial cytochrome b gene of 120 Rock Buntings from Germany, and a few birds from Switzerland, France and the Iberian Peninsula. In Figure 4, a phylogeny reconstruction by Maximum Likelihood (ML) is illustrated. All samples cluster in a single clade as a sister to the Cirl Bunting, which often occurs in the same regions. We have also obtained cyt b sequences for over 60 Cirl Buntings. All Rock Buntings and all Cirl Bunting clustered in separated clades. Thus, no apparent hybrids between both species could be detected, i.e. all samples of *E. cia* and *E. cirlus* clustered as separate groups . Cyt b sequences show a low level of genetic diversity and no apparent geographical structure.

**Figure 4:**
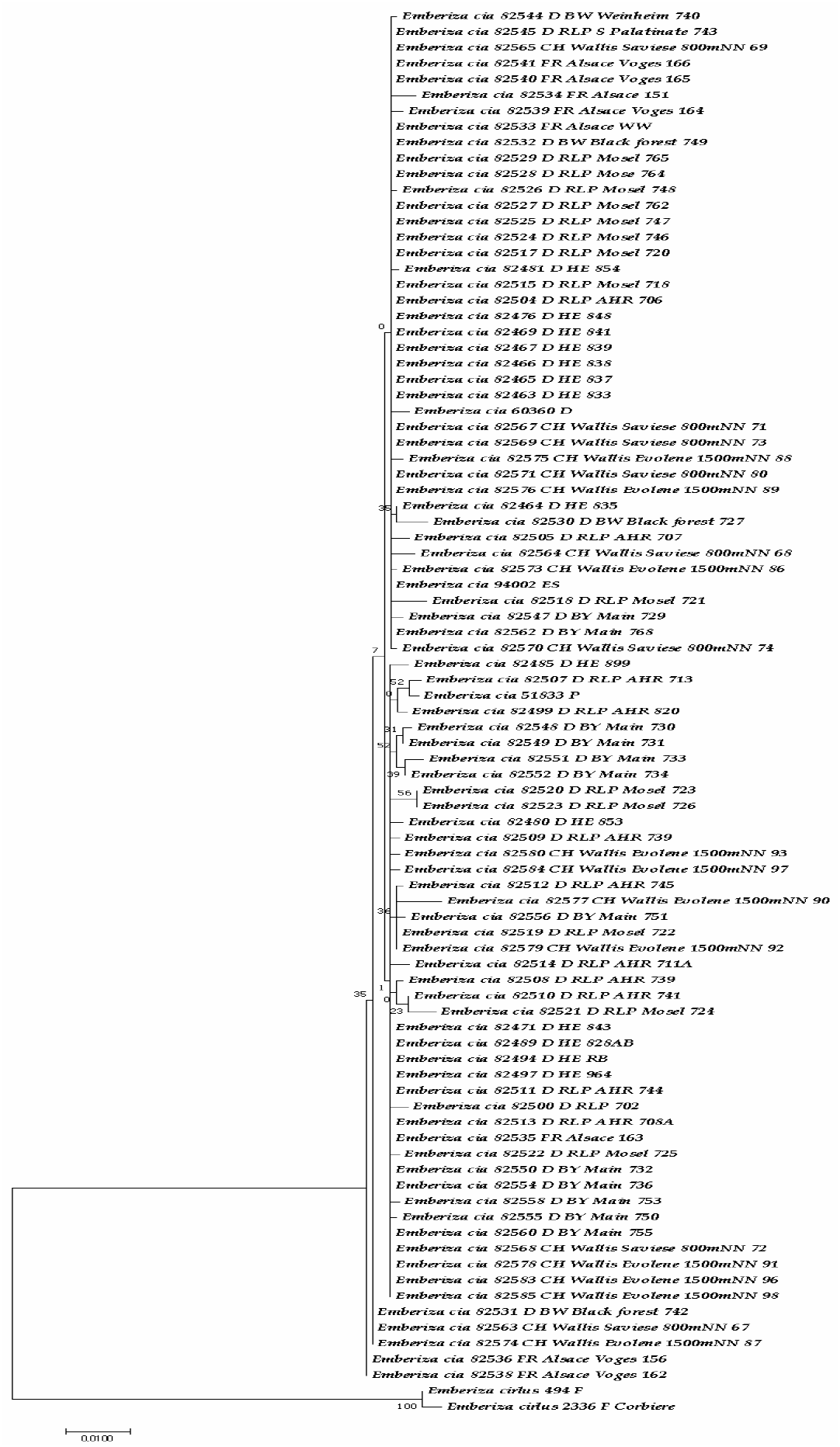
Molecular Phylogenetic analysis of *E. cia* individuals by the Maximum Likelihood method. The evolutionary history was inferred by using the Maximum Likelihood method based on the General Time Reversible model (Kimura 1980). The tree with the highest log likelihood (-2540,25) is shown. The percentage of trees in which the associated taxa clustered together is shown next to the branches. Initial tree(s) for the heuristic search were obtained automatically by applying Neighbor-Join and BioNJ algorithms to a matrix of pairwise distances estimated using the Maximum Composite Likelihood (MCL) approach, and then selecting the topology with superior log likelihood value. A discrete Gamma distribution was used to model evolutionary rate differences among sites (5 categories (+G, parameter = 0,2960)). The rate variation model allowed for some sites to be evolutionarily invariable ([+I], 0,00% sites). The tree is drawn to scale, with branch lengths measured in the number of substitutions per site. The analysis involved 89 nucleotide sequences. Codon positions included were 1st+2nd+3rd+Noncoding. There were a total of 1143 positions in the final dataset. Evolutionary analyses were conducted in MEGA 11 (Tamura et al. 2021). Numbers at branches refer to Bootstrap values (in %).

In order to better understand the genetic diversity, the haplotypes were identified in the data set (Tab. 4; Supplement Tab. S2) (haplotype diversity HD: 0.7158). The relationships between haplotypes were determined by Network analysis (Figure 5). Two major haplotypes exist from which the other haplotypes derive, often separated by one or two nucleotide differences (Figure 5).

**Figure 5:**
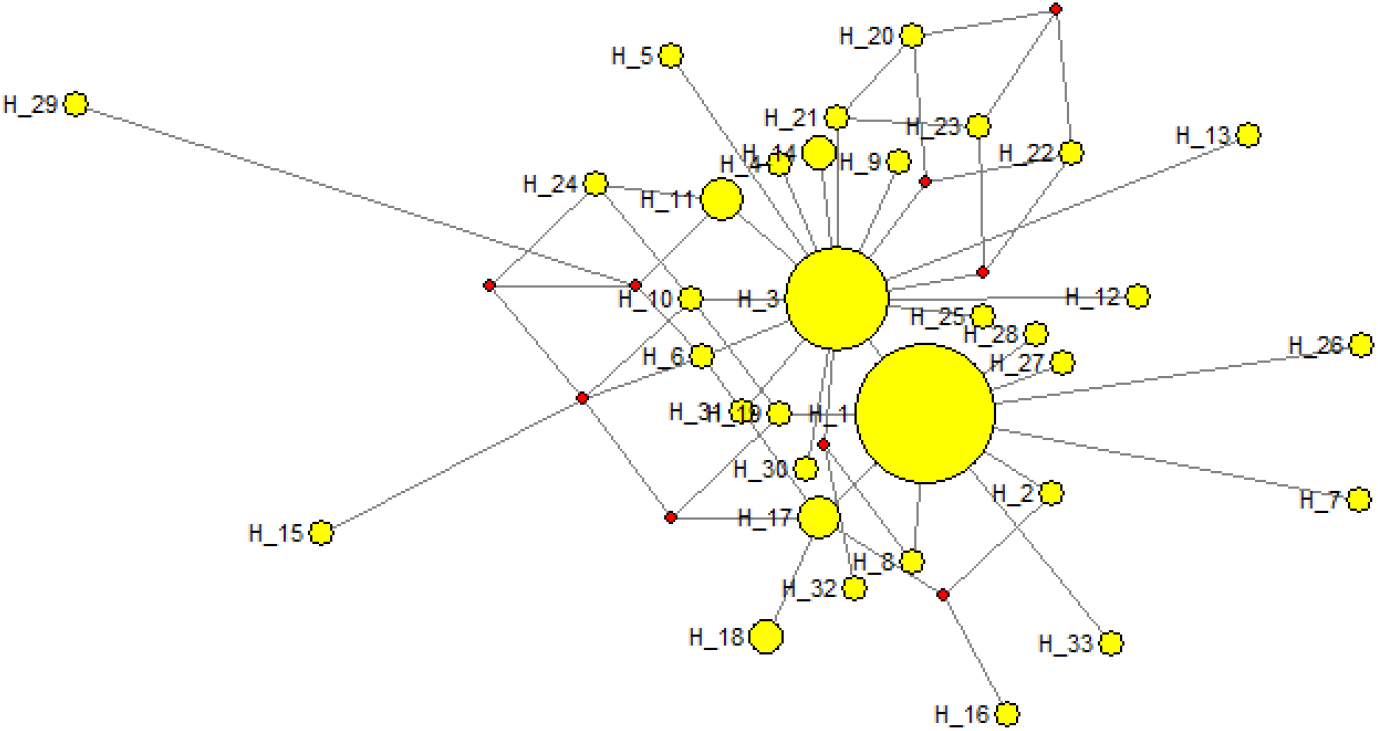
Haplotype network of cytochrome b of Rock buntings from Germany, France and Iberian Peninsula. Haplotype are in accordance to Table 4 and Table S2.

The most common haplotype H1 is found in birds from the rivers Moselle, Ahr, Main, Rhine, Black Forest, Alsace, Canton of Valais and Spain, whereas the second most common haplotype 2 occurred at Ahr, Moselle, Main, Rhine, and Canton of Valais. This means, that the haplotypes are not restricted to certain areas but occur as a mixture in the metapopulation (panmixia). The data indicate gene flow and connectivity within the metapopulation.

**Table 4:**
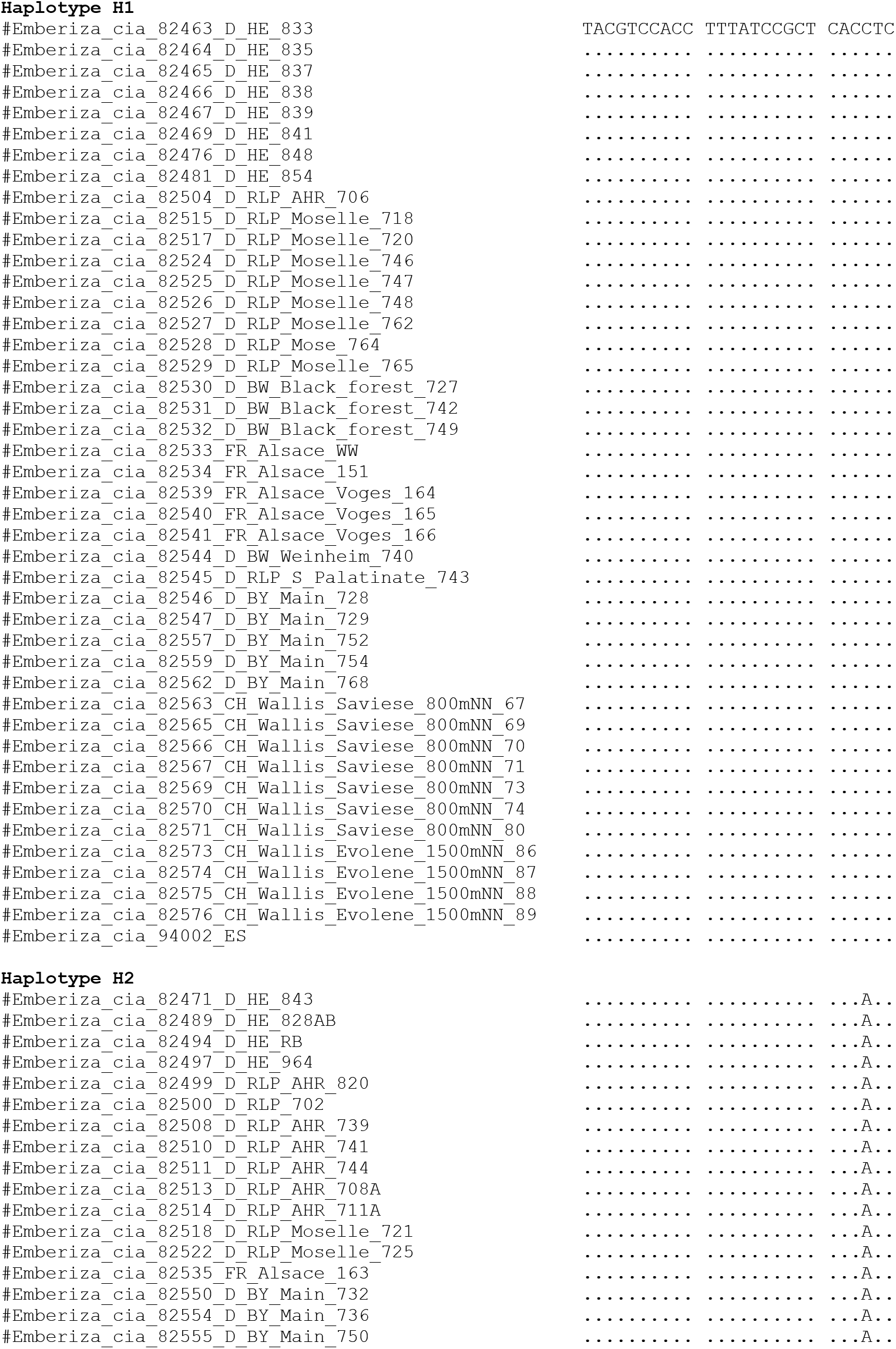
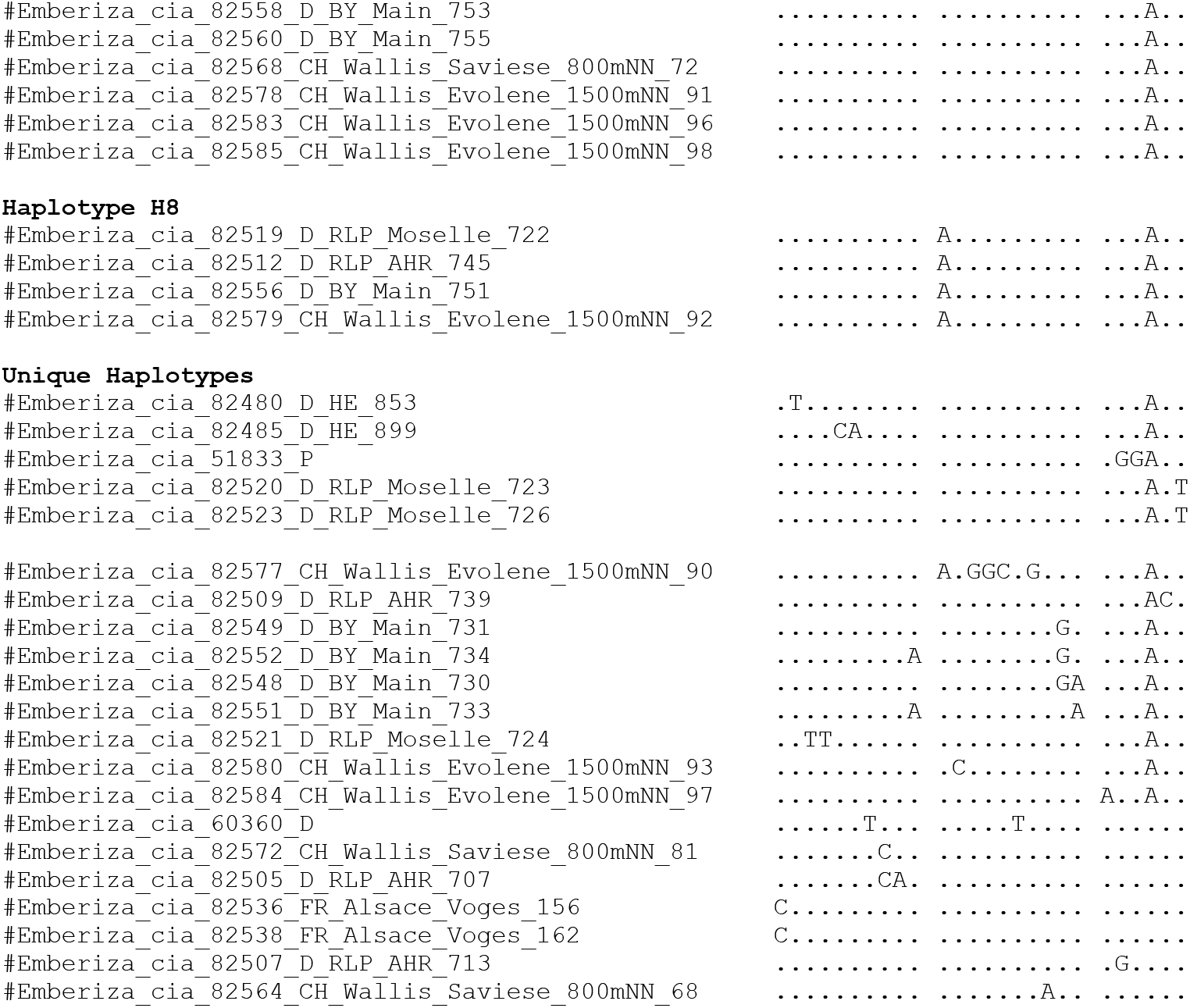
Attribution of nucleotide sequences of the mitochondrial cytochrome b gene of each sequenced bird to haplotypes. Variable sites in the cytochrome b sequence data set (without sites with missing data)

Such a panmixia is typical for many birds of the Palaearctic, such as Hoopoe, Bee-eater or Red-backed Shrike which have colonized mots part of Europe after the last glaciation (Wang et al., 2017; Carneiro et al., 2019; Parau et al., 2019; Parau and Wink 2021). Before, they had retracted to refuge areas in the Mediterranean, where lineages came together and mixed (Parau and Wink 2021). Many European migrant birds show two main haplotypes surrounded by small haplotype satellites, indicating that these populations may have derived from lineages in the eastern and western range of the Palaearctic, where were separated before the Miocene when glaciation cycles started (Parau and Wink 2021).

#### 3.2.2 Microsatellite analysis

As a nuclear marker we studied the allele distribution in 12 polymorphic microsatellite loci which were mostly in Hardy-Weinberg and linkage equilibrium (Table S3). Large allele dropout and null alleles were not detected. Private alleles were between 5 (Alsace, Ahr) and 17 (Rhine); Samples from Moselle and Main did not show private alleles. The total number of alleles to all 122 samples per locus was in the range 1 and 16 alleles at the 12 loci (Table 5).

**Table 5:** Number of alleles for each microsatellite locus and population.

|  | **Bmv** | **BWs** | **CH** | **F** | **Fe** | **HEr** | **RLPaw** | **RLPm** |
| --- | --- | --- | --- | --- | --- | --- | --- | --- |
| Embc-4 | 5 | 4 | 7 | 2 | 4 | 9 | 5 | 5 |
| Embc-5 | 6 | 4 | 7 | 4 | 8 | 14 | 12 | 7 |
| Embc-28 | 10 | 4 | 8 | 3 | 7 | 10 | 6 | 8 |
| Embc-13 | 7 | 1 | 8 | 2 | 4 | 8 | 5 | 5 |
| Embc-29 | 6 | 3 | 11 | 4 | 4 | 16 | 11 | 7 |
| Embc-14 | 7 | 3 | 11 | 4 | 5 | 12 | 7 | 6 |
| Embc-16 | 7 | 2 | 8 | 3 | 4 | 8 | 5 | 8 |
| Embc-22 | 6 | 3 | 8 | 3 | 4 | 12 | 7 | 5 |
| Embc-21 | 4 | 1 | 8 | 3 | 5 | 5 | 6 | 6 |
| Embc-19 | 4 | 3 | 7 | 6 | 6 | 8 | 5 | 4 |
| Embc-8 | 5 | 3 | 10 | 4 | 5 | 6 | 4 | 3 |
| Embc-30 | 4 | 2 | 6 | 4 | 4 | 5 | 5 | 5 |

Abbreviation of populations: CH= Wallis, HEr = middle Rhine, RLPm = Moselle, RLPaw = Ahr, BWs = Black Forest; Bms = Main; Fe= Alsace; F = France, Spain, Portugal

The average allelic richness (A_r_) varied from 4.4 and 5.9. The average gene diversity (H_s_) of all loci was 0.717 (range 0.456-0.867), whereas the overall gene diversity (H_T_) range was 0.458-0.885 (Table 6). The observed heterozygosity (H_o_) differed between 0.69 and 0.78 and the expected heterozygosity (H_e_) between 0.69 to 0.78 (Table 6). For the complete data set, He and Ho had a normal distribution and differed significantly (p= 0.0121).

**Table 6:**
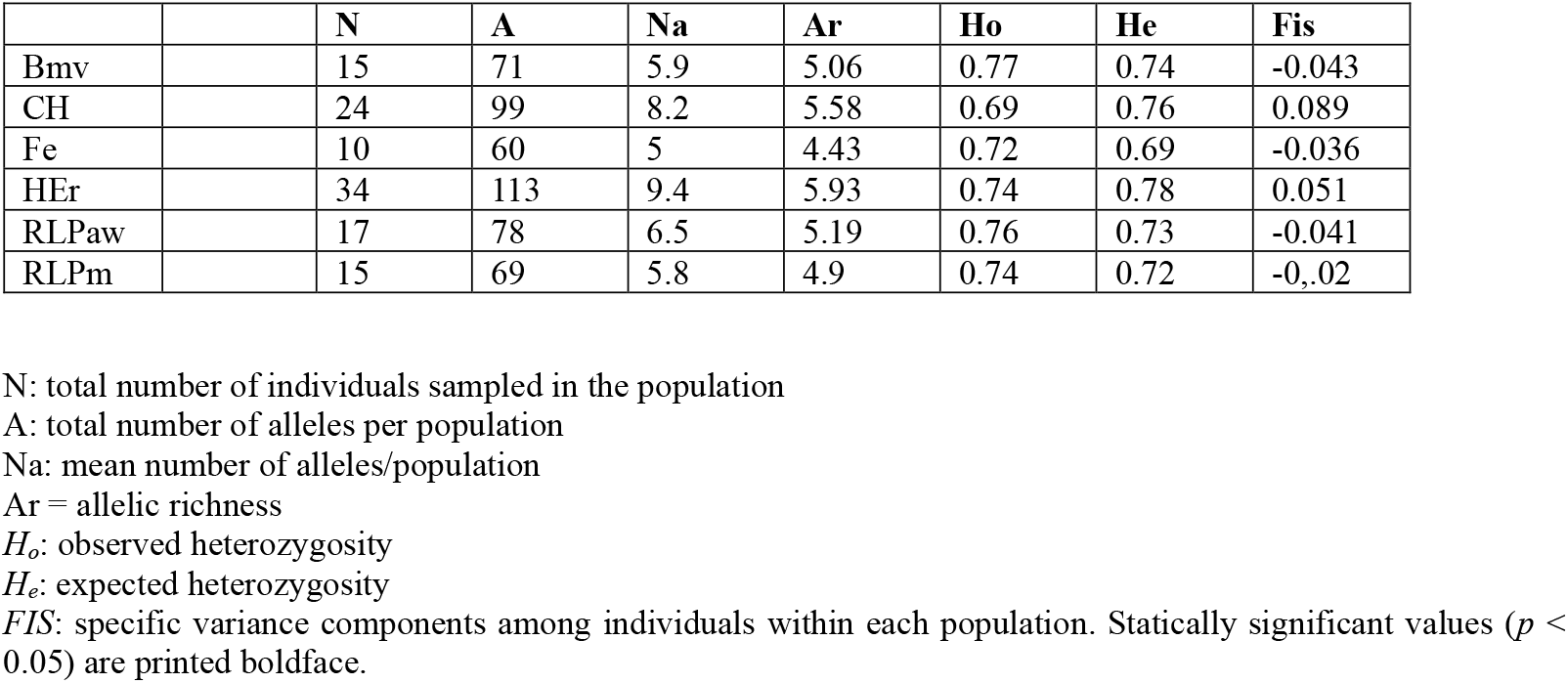
Summary statistics for the 12 microsatellite loci and six populations.

|  |  | **N** | **A** | **Na** | **Ar** | **Ho** | **He** | **Fis** |
| --- | --- | --- | --- | --- | --- | --- | --- | --- |
| Bmv |  | 15 | 71 | 5.9 | 5.06 | 0.77 | 0.74 | -0.043 |
| CH |  | 24 | 99 | 8.2 | 5.58 | 0.69 | 0.76 | 0.089 |
| Fe |  | 10 | 60 | 5 | 4.43 | 0.72 | 0.69 | -0.036 |
| HEr |  | 34 | 113 | 9.4 | 5.93 | 0.74 | 0.78 | 0.051 |
| RLPaw |  | 17 | 78 | 6.5 | 5.19 | 0.76 | 0.73 | -0.041 |
| RLPm |  | 15 | 69 | 5.8 | 4.9 | 0.74 | 0.72 | -0.02 |

N: total number of individuals sampled in the population
A: total number of alleles per population
Na: mean number of alleles/population
Ar = allelic richness
$H_o$ : observed heterozygosity
$H_e$ : expected heterozygosity
*FIS*: specific variance components among individuals within each population. Statistically significant values ( $p < 0.05$ ) are printed boldface.

Data were analysed with the program STRUCTURE in order to detect potential geographic signals within the metapopulation (Figure 6).

**Figure 6:**
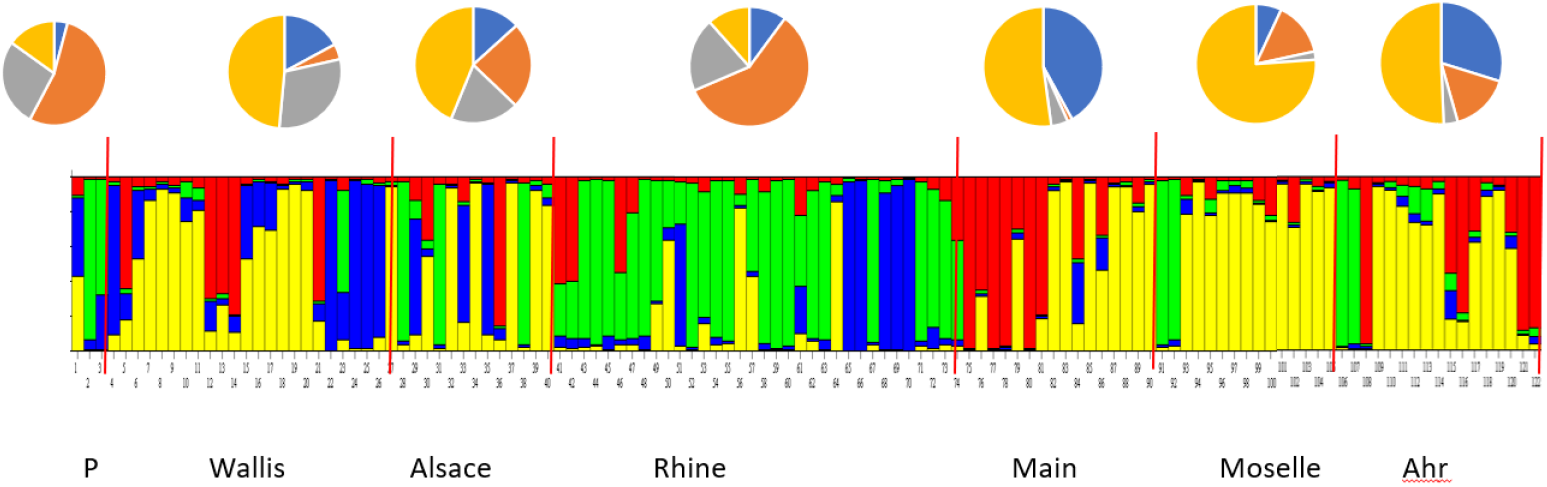
Graphic representation of a Bayesian Cluster Analysis with STRUCTURE, based on 12 microsatellite loci, represented with a K of 4. Conditions: Admixture, frequence correlated. The names of the populations are provided below the graphics. Each sample is represented as a column; the colours indicate the genetic clusters to which the alleles may belong. P= samples from Portugal, Spain, Southern France.

Bayesian clustering analyses using the Δ*K* method (Evanno et al., 2005) suggested that individuals can be attributed to four genetic clusters: Birds from the rivers Ahr, Moselle, Main, Alsace and partly Canton of Valais show a similar STR pattern, which is mostly different from Rock Buntings from southern France and the Iberian Peninsula. The Rock Buntings from the Rhine valley area express a different pattern, suggesting that they came from a different lineage. The STR data do not correlate with the cyt b haplotype network, suggesting that the divergence of mtDNA must be older than the STR pattern.

In another set of analysis, we employed a Discriminant Analysis of Principal Components (DAPC) based on 12 microsatellites (Figure 7). Again, subpopulations share many alleles and do not cluster in separate groups. Thus, also DAPC supports the results of our STRUCTURE and cyt b network analyses, suggesting an ongoing connectivity and genetic exchange between subpopulations.

**Figure 7:**
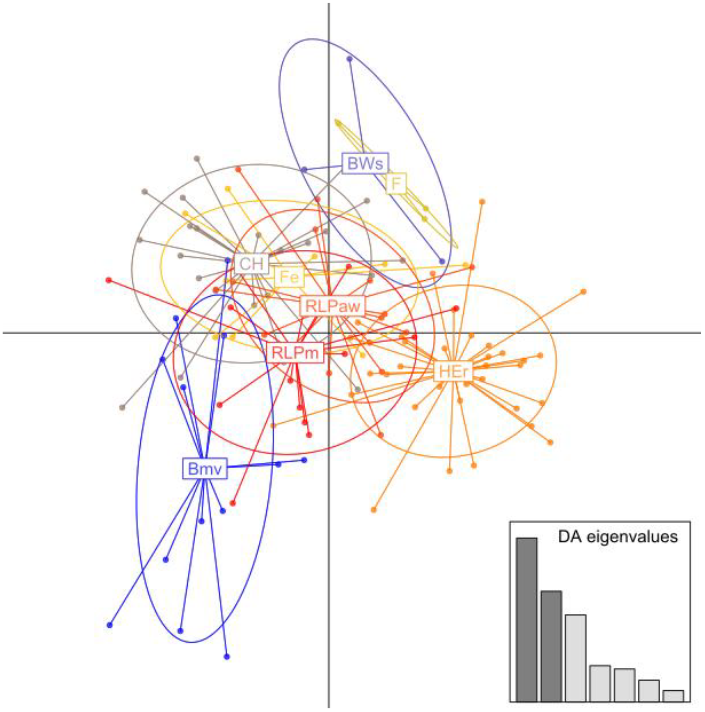
Discriminant Analysis of Principal Components (DAPC) based on 12 microsatellite loci of the Rock buntings. The dots correspond to the individuals, while the ellipses represent the cluster. CH= Wallis, Her = middle Rhine, RLP m = Moselle, RLP aw = Ahr, BWs = Black Forest; Bms = Main; Fe= Alsace; F = France, Spain, Portugal

## 4. Conclusions

All occurrences of the Rock Bunting in Germany and in neighbouring countries form spatially discrete and fragmented local populations. This is apparently due to the special habitat demands of Rock Buntings. Those habitats are characterized by steep rocky hillsides with xeric grassland. Two habitat differences are visible: Rocky, steep and south directed mountain hillsides (North- and South-Black Forest and Vosges up till over 1000 m, Canton of Valais till over 2000 m) and hillslopes along riverbanks (xerotherm steep habitats including wine terraces in about 200 m). The observed spatially discrete populations in these habitats show characteristics of subpopulations which potentially form two metapopulations, which are linked by more or less dispersal events. (Figure 2). The very localized subpopulations provide evidence for a restricted gene exchange. This assumption was supported by the observation that subpopulations of the Black Forest and the Palatinate became extinct and were not recolonized.

The genetic results provide no information of geographic differentiation so that the analysed subpopulation appear to be linked by more or less frequent dispersal events. The Rock Buntings from the area around Rüdesheim express a different pattern, suggesting that they came from a different lineage. Although a certain degree of genetic structure is apparent, we conclude that Rock Buntings of Central Europe form a single metapopulation with some degree of gene flow between the isolated subpopulations.

## Acknowledgements

We thank the following institutions for support of these investigations, for conferring the admission for bird ringing and blood sampling, and for special permits to use the car in restricted areas and to collect blood samples: Vogelwarte Helgoland, Institut für Vogelforschung, Wilhelmshaven und der Deutschen Ornithologen-Gesellschaft (DO-G) für eine Forschungsbeihilfe (Prof. Dr. Franz Bairlein); Vogelwarte Radolfzell, Max-Planck-Institut für Ornithologie Radolfzell (Dr. Wolfgang Fiedler); Staatliche Vogelschutzwarte für Hessen, Rheinland-Pfalz und Saarland, Institut für angewandte Vogelkunde (Gerd Bauschmann); Schweizerische Vogelwarte Sempach (Hannes v. Hirschheydt, Prof. Dr. Lukas Jenny); Muséum National d’Histoire Naturelle, Paris, (Dr. Olivier Dehorter); Prefecture du Haut-Rhin, Direction régional de Environement Alsace, Colmar; Untere Naturschutzbehörde Rhein-Taunuskreis, Bad Schwalbach, Hessen (Dr. Michael Berger); Struktur- und Genehmigungsdirektion Nord, Koblenz, Rheinland-Pfalz (Manfred Braun); Struktur- und Genehmigungsdirektion Süd, Neustadt/Weinstrasse, Rheinland-Pfalz (Thomas Schlindwein); Regierungspräsidium Freiburg, Baden-Württemberg, Abteilung Umwelt (Referat 56, Uwe Kerkhof); Bayerisches Landesamt für Umwelt (LfU), Vogelschutzwarte, Garmisch-Partenkirchen (Günther von Lossow); Regierung von Unterfranken, Höhere Naturschutzbehörde, Würzburg (Peter Krämer); Jean-Jacques Pfeffer, Linthal, Haut-Rhin (Alsace), Marc und Cleo Weibel, Linthal-Remspach, Haut-Rhin (Alsace) Dipl.-Biol. Florian Straub, Albert-Ludwigs Universität Freiburg Météo France provided the der climate data (Vosges); Alan Slusarenko for assistance and help for the English version

## Notes

### Competing Interest Statement

The authors have declared no competing interest.

## Literature

Arctander P. (1988): Comparative studies of avian DNA by restriction fragment length polymorphism analysis: Convenient procedures based on blood samples from live birds. Journal for Ornithology 129, 205–216.

Bosselmann J. (2008): Zippammer-Beobachtungen (Emberiza cia) 2005-2008 in Rheinland-Pfalz, Bestandsschätzungen. Pflanzen und Tiere in Rheinland-Pfalz 18, 152-155, Mayen.

Carneiro de Melo Moura C., A. Bastian, H-V. Bastian, E. Wang, X. Wang and M. Wink (2019): Pliocene origin, ice ages and postglacial population expansion have influenced a panmictic phylogeography of the European Bee-eater Merops apiaster. Diversity 2019, 11, 12; doi:10.3390/d11010012.

Deuschle J., F. Straub, D. Kratzer, I. Schuphan, U. Dorka and A. Plank (2010): Natura 2000 Managementplan „Südschwarzwald”, MaP-Bearbeitung der Zippammer (Emberiza cia L.) in Vogelschutzgebieten Baden-Württembergs (MaP-Gebiete 2009-1010), Teilbeitrag für das Vogelschutzgebiet 8441-441Südschwarzwald, Landesamt für Umwelt, Messungen und Naturschutz, Baden-Württemberg (LUBW), Karlsruhe.

Dietzen C., H.-G. Folz, T. Grunwald, P. Keller, A. Kunz, M. Niehuis, M. Schäf, M. Schmolz and M. Wagner (2017): Die Vogelwelt in Rheinland-Pfalz. Band 4.2 Singvögel (Passeriformes). Fauna und Flora in Rheinland-Pfalz, Beiheft 49, Landau.

Dorka U. (2010): in Deuschle J., F. Straub, D. Kratzer, I. Schuphan, U. Dorka and A. Plank (2010): Natura 2000.

Evanno G., S. Regnaut, J. Goudet (2005): Detecting the number of clusters of individuals using the software STRUCTURE: a simulation study. Molecular Ecology 14, 2611–2620.

Fuchs F.-J. and T. Macke (2002): Verbreitung der Zippammer im Ahrtal. Ergebnis der Revierkartierungen 1997 und 1999.- Fauna und Flora in Rheinland-Pfalz, Beiheft 27, 263-266. Landau.

Glutz von Blotzheim U.N. and K. M. Bauer (1997): Handbuch der Vögel Mitteleuropas. Band 14, Passeriformes (5. Teil). Aula,Wiesbaden.

Groh G. (1988): Zur Biologie der Zippammer (Emberiza cia cia L.) im Pfälzerwald. Mitt. Pollichia 75, 261–287.

Guo SW, and EA Thompson (1992): Performing the exact test of Hardy-Weinberg proportion for multiple alleles. Biometrics, 48, 361–372.

Hanski I. and O. Ovaskainen (2003): Metapopulation theory for fragmented landscapes, Theoretical Population Biology 64, 119–127.

Jombart T., S. Devillard and F. Balloux (2010): Discriminant analysis of principal components: a new method for the analysis of genetically structured populations. BMC Genetics 11, 94. doi: 10.1186/1471-2156-11-94.

Keller V., S Herrando el al. (2020). European Breeding Bird Atlas 2. European Bird Census Council and Lynx Edicions, Barcelona.

Keusch P. (1991): Vergleichende Studie zu Brutbiologie, Jungenentwicklung, Bruterfolg und Populationsökologie von Ortolan (Emberiza hortulana) und Zippammer (Emberiza cia) im Alpenraum, mit besonderer Berücksichtigung des unterschiedlichen Zugverhaltens. Diss. Univ. Bern.

Kimura M. (1980): A simple method for estimating evolutionary rate of base substitutions through comparative studies of nucleotide sequences. Journal of Molecular Evolution 16,111–120.

Lemoine N, H-G Bauer, M. Peintinger and K. Böhning-Gaese (2007): Effects of climate and land-use change on species abundance in a central European bird community. Conservation Biology 21, 495–503.

Levins R. (1970): Extinction. In: Some Mathematical Questions in Biology (ed. M. Gerstenhaber), pp. 77 – 107. Providence, RI: The American Mathematical Society

Maumary L., L. Vallotton and P. Knaus (2007): Die Vögel der Schweiz. Schweizerische Vogelwarte, Sempach, und Nos Oiseaux, Montmollin.

Nei M. and S. Kumar (2000): Molecular Evolution and Phylogenetics. Oxford University Press, New York.

Pârâu LG., R. Frias Soler and M. Wink (2019): High genetic diversity among breeding Red-backed Shrikes Lanius collurio in the Western Palearctic. Diversity 11, 31; doi: 10.3390/d11030031.

Pârâu L., and M. Wink (2021): Common patterns in the molecular phylogeography of Western Palearctic birds: A comprehensive review. Journal of Ornithology. 162, 937–959 10.1007/s10336-021-01893

Pfeffer J-J. and Gilot F. (2002): Statut du Bruant fou (Emberiza cia) dans les Vosges Haut-Rhinoises. Ciconia 26, 65–74.

Pritchard JK., M. Stephens and P. Donnelly (2000): Inference of population structure using multilocus genotype data. Genetics 155, 945–959.

R Core Team (2022): R: A language and environment for statistical computing. R Foundation for Statistical Computing, Vienna, Austria. https://www.R-project.org/.

Rozas J., A. Ferrer-Mata, J.C. Sánchez-DelBarrio, S. Guirao-Rico, P. Librado, S.E. Ramos-Onsins, and A. Sáchez-Gracia (2017): DnaSP v6: DNA Sequence Polymorphism Analysis of Large Datasets. Molecular Biology and Evolution 34, 3299–3302.

Schlotmann F. and E. Dietrich (2012): Die Avifauna der Weinbaugebiete in Rheinland-Pfalz.- Fauna and Flora 12, 629–702.

Schmid H., R. Luder, B. Naef-Daenzer, R. Graf and N. Zbinden (1998): Schweizer Brutvogelatlas. Verbreitung der Brutvögel in der Schweiz und im Fürstentum Liechtenstein 1993–1996. Schweizerische Vogelwarte, Sempach.

Schuphan I. (1972): Zur Biologie und Populationsdynamik der Zippammer (Emberiza c. cia L.). Diplomarbeit. Johannes Gutenberg-Univ. Mainz. http://www.hgon.de/service/downloads/

Schuphan I. (2007): Langfristige Einflüsse von Pflegemaßnahmen, Flurbereinigung und Klimaerwärmung auf eine farbig beringte Teilpopulation der Zippammer Emberiza cia am Mittelrhein. Vogelwarte 45, 299–300.

Schuphan I. (2011a): Habitat-Strukturen und populationsdynamische Parameter einer Population der Zippammer (Emberiza cia): Nutzbare Basisdaten für zukünftige Zippammer–Managementpläne. Vogelwarte 49: 65–74.

Schuphan I. (2011b): Die Zippammer (Emberiza cia) eine große Klimaunterschiede ertragende Vogelart. Vogelwarte 49, 129–136.

Schuphan I. (2011c): Bestand der Zippammer Emberiza cia 2011am Kallmuth bei Homburg am Main und am Main zwischen Karlstadt und Veitshöchheim. Ornithologischer Anzeiger 50, 133–141.

Schuphan I. (2017): Zippammer Emberiza cia Linnaeus 1766. In Dietzen C. et al.: Die Vogelwelt in Rheinland-Pfalz. Band 4.2 Singvögel (Passeriformes). – Fauna und Flora in Rheinland-Pfalz, Beiheft 49, 1021–1038. Landau.

Schuphan I, and B. Flehmig (2022): Zippammer – Emberiza cia Bestand im Unteren Rheingau: Dramatischer und fortlaufender Rückgang seit Beginn der Flurbereinigung vor 60 Jahren. Vogelwarte 60, 51–60.

Schuphan I. and F. Grimm (2012): Die Zippammer (Emberiza cia) in der Südpfalz. Fauna und Flora in Rheinland-Pfalz 12, 703–712.

Schuphan I. and M. Wink (2016): The Rock Bunting Emberiza cia in Southwest Switzerland: From hot rock steppes to rough high mountains. Ornithologischer Beobachter 113: 299–308.

Segelbacher G. and I. Schuphan (2014): Isolation of 13 tetranucleotide microsatellite loci in the Rock Bunting (Emberiza cia). Conservation Genetics Resources, DOI 10.1007/s12686-014-0149-0.

Stein F-J (2011): In: Ausgewählte Ergebnisse der Bestandsaufnahmen genauer erfasster Arten für das Jahr 2010. Irrgeister, Naturmagazin HSK e.V. 28: 24.

Storch V., U. Welsch, M. Wink (2013): Evolutionsbiologie. 3. edition; Springer, Heidelberg.

Tamura K., G. Stecher G, and S. Kumar (2021): MEGA11: Molecular Evolutionary Genetics Analysis version 11. Molecular Biology and Evolution 38, 3022–3027.

Wang E., R. E. Van Wijk, M. S. Braun and M. Wink (2017): Gene flow and genetic drift contribute to high genetic diversity with low phylogeographical structure in European Hoopoes (Upupa epops). Molecular Phylogeny and Evolution,113, 113–125.

