## Supplementary material for "Rock Buntings in Central Europe: Phylogeographic connectivity and a current inventory in the Upper Middle Rhine valley": Table 1 to 3

### Supplements:

### Table S1: Details of Rock Bunting monitoring (by IS)

| Rock Bunting-occurrence | Visiting days | | | | | | | | | | | |
|---|---|---|---|---|---|---|---|---|---|---|---|---|
| | 2007 | 2008 | 2009 | 2010 | 2011 | 2012 | 2013 | 2014 | 2015 | 2016 | 2017 | 2018 |
| Region Brillon-NRW | | | | | | 1 | | | | | | |
| River Ahr | 2 | 2 | 2 | 1 | 1 | - | | | | | | |
| River Moselle | 3 | 4 | 5 | 4 | 2 | 1 | | | | | | |
| River Middle Rhine (federal state Hesse) | 4 | 4 | 6 | 5 | 6 | 6 | 9 | 8 | 8 | 11 | 10 | 9 |
| River Nahe | 2 | | 1 | | 1 | - | | | | | | |
| River Main | - | 3 | 3 | - | 3 | 1 | | | | | | |
| Palatinate | - | 1 | 1 | 1 | 1 | 3 | 3 | 3 | 3 | | | 2 |
| Region Bergstraße | - | - | 3 | 1 | - | - | | | | 1 | 1 | |
| South-Black Forest | | 3 | 4 | 3 | 3 | 3 | | | | | | |
| Vosgeses | | 3 | 6 | 3 | 5 | 4 | 6 | | | | | |
| Lake Constance | | | 1 | 1 | 1 | - | | | | | | |
| Canton of Valais, VS | | | | | | 12 | | 6 | 3 | | | |

### Table S2: Haplotype statistics

| **Haplotype** | **Frequency** | **Origin** |
|---|---|---|
| Hap_1: | 43 | Emberiza_cia_82463_D_HE_833<br>Emberiza_cia_82464_D_HE_835<br>Emberiza_cia_82465_D_HE_837<br>Emberiza_cia_82466_D_HE_838<br>Emberiza_cia_82467_D_HE_839<br>Emberiza_cia_82469_D_HE_841<br>Emberiza_cia_82476_D_HE_848<br>Emberiza_cia_82481_D_HE_854 |

| | | |
|---|---|---|
| | | Emberiza_cia_82504_D_RLP_AHR_706<br>Emberiza_cia_82515_D_RLP_Moselle_718<br>Emberiza_cia_82517_D_RLP_Moselle_720<br>Emberiza_cia_82524_D_RLP_Moselle_746<br>Emberiza_cia_82525_D_RLP_Moselle_747<br>Emberiza_cia_82526_D_RLP_Moselle_748<br>Emberiza_cia_82527_D_RLP_Moselle_762<br>Emberiza_cia_82528_D_RLP_Mose_764<br>Emberiza_cia_82529_D_RLP_Moselle_765<br>Emberiza_cia_82530_D_BW_Black_forest_727<br>Emberiza_cia_82531_D_BW_Black_forest_742<br>Emberiza_cia_82532_D_BW_Black_forest_749<br>Emberiza_cia_82533_FR_Alsace_WW<br>Emberiza_cia_82534_FR_Alsace_151<br>Emberiza_cia_82539_FR_Alsace_Voges_164<br>Emberiza_cia_82540_FR_Alsace_Voges_165<br>Emberiza_cia_82541_FR_Alsace_Voges_166<br>Emberiza_cia_82544_D_BW_Weinheim_740<br>Emberiza_cia_82545_D_RLP_S_Palatinate_743<br>Emberiza_cia_82546_D_BY_Main_728<br>Emberiza_cia_82547_D_BY_Main_729<br>Emberiza_cia_82557_D_BY_Main_752<br>Emberiza_cia_82559_D_BY_Main_754<br>Emberiza_cia_82562_D_BY_Main_768<br>Emberiza_cia_82563_CH_Wallis_Saviese_800mNN_67<br>Emberiza_cia_82565_CH_Wallis_Saviese_800mNN_69<br>Emberiza_cia_82566_CH_Wallis_Saviese_800mNN_70<br>Emberiza_cia_82567_CH_Wallis_Saviese_800mNN_71<br>Emberiza_cia_82569_CH_Wallis_Saviese_800mNN_73<br>Emberiza_cia_82570_CH_Wallis_Saviese_800mNN_74<br>Emberiza_cia_82571_CH_Wallis_Saviese_800mNN_80<br>Emberiza_cia_82573_CH_Wallis_Evolene_1500mNN_86<br>Emberiza_cia_82574_CH_Wallis_Evolene_1500mNN_87<br>Emberiza_cia_82575_CH_Wallis_Evolene_1500mNN_88<br>Emberiza_cia_82576_CH_Wallis_Evolene_1500mNN_89 |
| Hap_2: | 23 | Emberiza_cia_82471_D_HE_843<br>Emberiza_cia_82489_D_HE_828AB<br>Emberiza_cia_82494_D_HE_RB<br>Emberiza_cia_82497_D_HE_964<br>Emberiza_cia_82499_D_RLP_AHR_820<br>Emberiza_cia_82500_D_RLP_702<br>Emberiza_cia_82508_D_RLP_AHR_739<br>Emberiza_cia_82510_D_RLP_AHR_741<br>Emberiza_cia_82511_D_RLP_AHR_744<br>Emberiza_cia_82513_D_RLP_AHR_708A<br>Emberiza_cia_82514_D_RLP_AHR_711A<br>Emberiza_cia_82518_D_RLP_Moselle_721<br>Emberiza_cia_82522_D_RLP_Moselle_725<br>Emberiza_cia_82535_FR_Alsace_163<br>Emberiza_cia_82550_D_BY_Main_732<br>Emberiza_cia_82554_D_BY_Main_736<br>Emberiza_cia_82555_D_BY_Main_750<br>Emberiza_cia_82558_D_BY_Main_753<br>Emberiza_cia_82560_D_BY_Main_755<br>Emberiza_cia_82568_CH_Wallis_Saviese_800mNN_72<br>Emberiza_cia_82578_CH_Wallis_Evolene_1500mNN_91<br>Emberiza_cia_82583_CH_Wallis_Evolene_1500mNN_96<br>Emberiza_cia_82585_CH_Wallis_Evolene_1500mNN_98 |
| Hap_3 | 1 | Emberiza_cia_82480_D_HE_853] |

| | | |
|---|---|---|
| Hap_4 | 1 | Emberiza_cia_82485_D_HE_899] |
| Hap_5 | 1 | Emberiza_cia_82505_D_RLP_AHR_707] |
| Hap_6 | 1 | Emberiza_cia_82507_D_RLP_AHR_713] |
| Hap_7 | 1 | Emberiza_cia_82509_D_RLP_AHR_739] |
| Hap_8 | 4 | Emberiza_cia_82512_D_RLP_AHR_745 Emberiza_cia_82519_D_RLP_Moselle_722 Emberiza_cia_82556_D_BY_Main_751 Emberiza_cia_82579_CH_Wallis_Evolene_1500mNN_92] |
| Hap_9 | 2 | Emberiza_cia_82520_D_RLP_Moselle_723 Emberiza_cia_82523_D_RLP_Moselle_726] |
| Hap_10 | 1 | Emberiza_cia_82521_D_RLP_Moselle_724] |
| Hap_11 | 2 | Emberiza_cia_82536_FR_Alsace_Voges_156 Emberiza_cia_82538_FR_Alsace_Voges_162] |
| Hap_12 | 1 | Emberiza_cia_82548_D_BY_Main_730] |
| Hap_13 | 1 | Emberiza_cia_82549_D_BY_Main_731] |
| Hap_14 | 1 | Emberiza_cia_82551_D_BY_Main_733] |
| Hap_15 | 1 | Emberiza_cia_82552_D_BY_Main_734] |
| Hap_16 | 1 | Emberiza_cia_82564_CH_Wallis_Saviese_800mNN_68] |
| Hap_17 | 1 | Emberiza_cia_82572_CH_Wallis_Saviese_800mNN_81] |
| Hap_18 | 1 | Emberiza_cia_82577_CH_Wallis_Evolene_1500mNN_90] |
| Hap_19 | 1 | Emberiza_cia_82580_CH_Wallis_Evolene_1500mNN_93] |
| Hap_20 | 1 | Emberiza_cia_82584_CH_Wallis_Evolene_1500mNN_97] |
| Hap_21 | 1 | Emberiza_cia_51833_P] |
| Hap_22 | 1 | Emberiza_cia_60360_D] |

**Table S3 HWE results (p values) for the subpopulations (ns= not significant; s = significant)**

| | **Bmv** | **BWs** | **CH** | **F** | **Fe** | **HEr** | **RLPaw** | **RLPm** |
|---|---|---|---|---|---|---|---|---|
| Embc-4 | ns | ns | ns | ns | ns | ns | ns | ns |
| Embc-5 | ns | ns | ns | ns | ns | ns | ns | ns |
| Embc-28 | ns | ns | s | ns | ns | ns | ns | ns |

| | | | | | | | | |
|---|---|---|---|---|---|---|---|---|
| Embc-13 | ns | ns | ns | ns | ns | ns | ns | ns |
| Embc-29 | ns | ns | ns | ns | ns | s | s | ns |
| Embc-14 | ns | ns | ns | ns | ns | ns | ns | ns |
| Embc-16 | ns | ns | ns | ns | ns | ns | ns | ns |
| Embc-22 | ns | ns | ns | ns | ns | ns | ns | ns |
| Embc-21 | ns | ns | ns | ns | ns | ns | ns | ns |
| Embc-19 | ns | ns | s | ns | ns | ns | ns | ns |
| Embc-8 | ns | ns | ns | ns | ns | ns | ns | ns |
| Embc-30 | ns | ns | ns | ns | ns | ns | ns | ns |
